# Synergy of Quorum Quenching Enzyme and Quorum Sensing Inhibitor in Inhibiting *P.aeruginosa* Quorum Sensing

**DOI:** 10.1101/182543

**Authors:** July Fong, Chaodong Zhang, Renliang Yang, Zhao Zhi Boo, Soon Keat Tan, Thomas E. Nielsen, Michael Givskov, Bin Wu, Haibin Su, Liang Yang

## Abstract

The threat of antibiotic resistant bacteria has called for alternative antimicrobial strategies that would mitigate the increase of classical resistance mechanism. Many bacteria employ quorum sensing (QS) to govern the production of virulence genes and formation of drug-resistance biofilms. Blocking QS mechanisms have proven to be a functional alternative to conventional antibiotic control of infections. The concepts of quorum sensing inhibitors (QSI) and quorum quenching enzymes (QQ) have been investigated separately. In this study however, we simulated the synergistic effect of QQ and QSI in blocking bacterial QS. This effect was validated by experiments using AiiA and G1 as QQ and QSI respectively on *Pseudomonas aeruginosa* LasR/I and RhlR/I QS circuits. The combination of a QQ and a QSI almost completely blocked the *P. aeruginosa* QS *las* and *rhl* system. Our findings provided a potential application strategy for bacterial QS disruption.

## Introduction

The emerging threat of antibiotic resistant bacterial pathogens has called for alternative strategies that could minimize the development of resistance mechanism. One such strategy is to interfere with the signaling pathways governing the social behaviors^1^. Microbial organisms exhibit social behaviors and communicate with each other through quorum sensing (QS)^2-4^. By synthesizing small signal molecules, they respond collectively to regulate expression of virulence factors, biofilm development, secondary metabolite production, interactions with host and other microbes in a population-density manner^5^. As QS is involved in bacterial behaviors in particular those causing diseases, targeting QS mechanisms has been put forward as an attractive approach to conventional infection control^1^.

Acylhomoserine lactone (AHL)-based QS signals are found in more than 70 bacterial species, in which many of them are pathogens^3,6^. In most cases, the structures of the AHLs are conserved with a homoserine lactone (HSL) ring connected to an acyl group with different chain length (n = 4-16)^5,7^. There are two AHL-mediated QS systems in the opportunistic pathogen *Pseudomonas aeruginosa*, which comprise the Lux homologues LasRI and RhlRI, respectively. LasRI and RhlRI function in the hierarchical manner in controlling the gene expression. LasI and RhlI are responsible for the synthesis of *N*-(3-oxododecanoyl) homoserine lactone (3-oxo-C12-HSL) and *N-*butanoylhomoserine lactone (C4-HSL) respectively, while the LasR and RhlR function as receptors for 3-oxoC12-HSL and C4-HSL and subsequently activate gene expression of QS target genes^8-10^. On top of that, there is also third signaling molecule, “pseudomonas quinolone signal” (PQS) which intertwined between the *las* and *rhl* systems^11^. QS defective *P. aeruginosa* mutants have much reduced virulence as compared to the wild-type strain and unable to establish infections in several animal models^1,12,13^.

The concept of QS disruption encompasses not just medicine and healthcare settings, but also membrane bioreactor, aquaculture and crop production^5,14^. It could be achieved by interfering with the QS signaling pathways (signal generator or receptor), or intercepting with the signal molecules (AHL)^15-17^. Enzymes that inactivate QS signals are called quorum quenchers (QQ), while chemicals that disrupt the QS pathways and reduce the expression of QS-controlled genes are called quorum sensing inhibitors (QSI)^5^. The first study on how quorum quenching enzyme could be used to control bacterial infections was demonstrated by Dong et al.^18^. The enzyme encoded by *aiiA* gene isolated from Gram-positive *Bacillus* species is capable of inactivating AHL signals through hydrolysis of the ester bond of the homoserine lactone ring and quench the QS signaling. It was proposed that the AHL-lactonase (AiiA) paralyses the QS signals and virulence factors production, hence allows the host defense mechanisms to halt and clear the bacterial infection^19^.

Mathematical modeling has been a useful tool to answer basic and conceptual research questions. In the last decade, mathematical modeling of QS has provided understanding to key components of the QS networks^20^. It has been used to examine *P. aeruginosa* LasR/I circuit and predict the biochemical switch between two steady states of system (low and high levels of signal perception) and QS response to colony size and cell density^21^. In another study, Magnus et al. included both LasR/I and RhlR/I circuits of *P. aeruginosa* in their model. Their results suggested Vfr increases the affinity between LasR-AHL dimer and LasR promoter, which was supported by experiments showing that Vfr was important at initial but not later stages of QS induction^22^. Goryachev et al. analyzed *Vibrio fischeri* QS and found that dimerization of LuxR-AHL is important for the stability of QS network^23^. Weber et al. considered individual cell heterogeneity and concluded that in *Vibrio fischeri* QS network, LuxR expression noise decreases autoinducer turning on threshold of single cell but slows down the population level QS induction^24^. Altogether, the models developed in these studies provide basic understanding of QS networks utilizing LuxIR regulatory system and its homologues, which are identified in many Gram-negative bacteria^25,26^.

In this study, we explored the concept of combining a QQ enzyme and a QSI compound to disrupt AHL signaling and signal reception capacities, and reduce the pathogenicity of *P. aeruginosa*. The QS network in *P. aeruginosa* is highly adaptable and capable of responding to the environmental stress conditions^27,28^, hence combinational therapy could provide multiple points of attack to increase bacteria coverage^29^. The two classes of QS disrupting agents have been studied independently, each with their own advantages and drawbacks. Small molecules as QSIs have well-known chemical structures, which in turn would allow structural activity and relationship (SAR) study and biological activities modification (ie. pharmacodynamics and pharmacokinetics properties). The molecules can also diffuse into the cells and target the receptors, in contrast to QQ enzymes that act extracellularly to degrade AHLs^30^. Because of their distinct molecular structures and functional mechanisms, it would be interesting to explore the possible synergistic effect of a QQ enzyme and a QSI molecule.

## Results

### Mathematical modeling shows synergistic effects between QQ enzyme and QSI on LasR/I circuit

In our study, we chose QQ enzyme AiiA in combination with G1, a small molecule of QSI which binds to LasR and RhlR^31^ as our models. When only a QQ enzyme (AiiA) was present, simulation results showed a distinct on and off states (Fig. 1A). η(QQ) represents the AHL degradation rate by the QQ enzyme. When the QQ enzyme concentration is low and η(QQ) is small, the stationary AHL concentration is high. However, when η(QQ) exceeds a threshold (2.8 × 10^-4^*s*^-1^), the stationary AHL concentration suddenly decreases to an insignificant value. Similar switching behaviors have been observed in the simulation response curves of QS components to population size^32^, cell volume fraction^21^, or to external AHL concentration^33^. Switching behaviors of QS networks have also been observed experimentally at individual cell level^34,35^.

**Figure 1.**
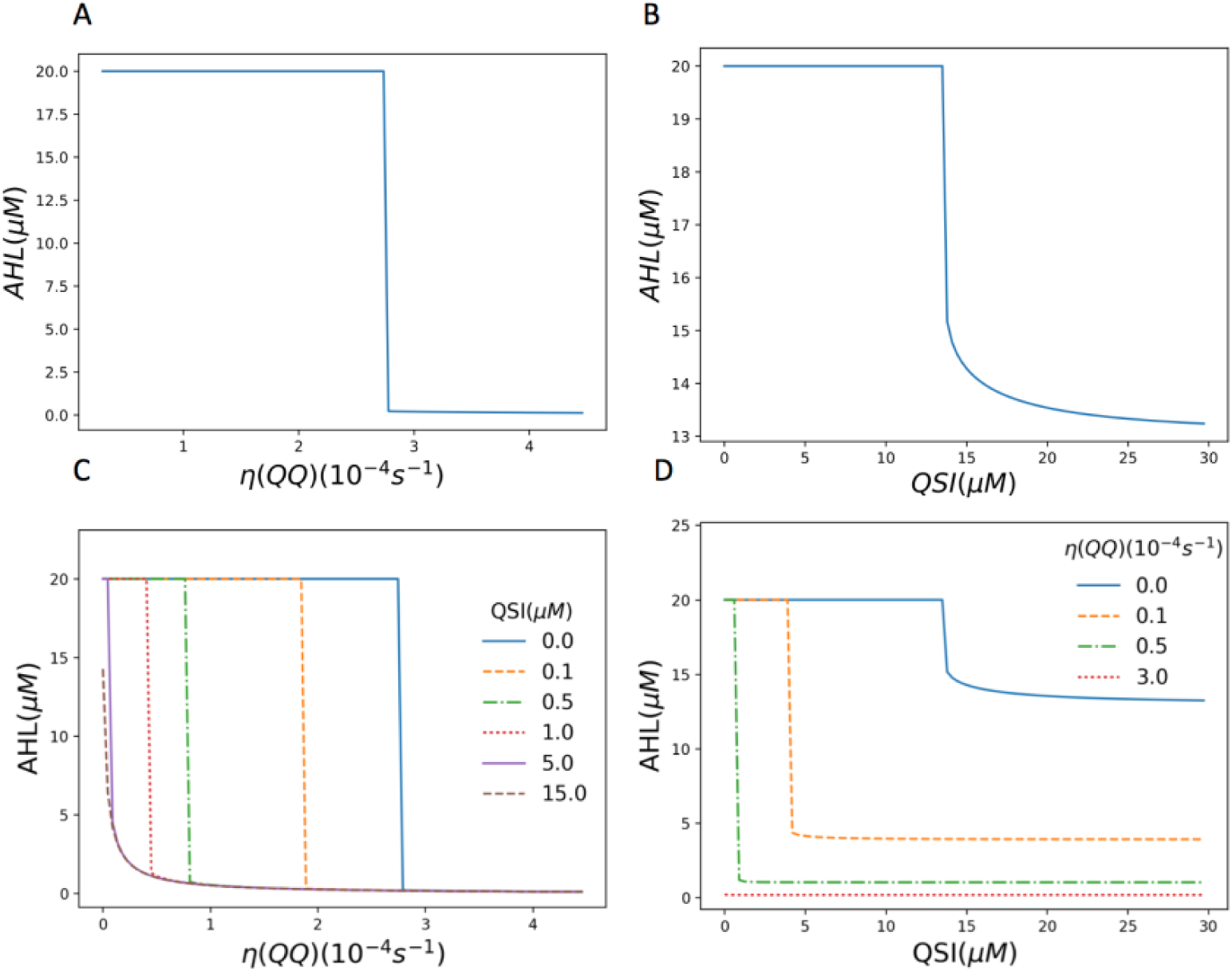
Simulation results of stationary AHL concentration to QQ and QSI. (A) η(QQ), (B) QSI, (C) η(QQ) at different QSI concentrations, and (D) QSI at different η(QQ) values.

When only the QSI was present, similar switching behavior was observed and shown in Fig. 1B. As the QSI and AHL bind to LasR competitively, the inhibiting effect of QSI is less efficient. Irreversible QSIs like halogenated furanones^36^ that induce degradation of the LasR receptor protein can inhibit QS more effectively (simulation data not shown). When combined, QQ and QSI can enhance the inhibiting effects of each other (Fig. 1C and D). 0.5 µM QSI alone has very little effect, but it can reduce the minimum QQ rate required to turn off QS up to 4 folds. Similarly, adding small amount of QQ (η(QQ) =0.5 × 10^-4^*s*^-1^) can reduce the minimum QSI concentration required to turn off QS up to 20 folds. The stationary AHL concentration decreased as compared to single treatment using QSI.

3D plot of stationary AHL to η(QQ) and QSI is shown in Fig. 2A. A clear boundary between QS on and off states was observed, which is shown in 3B. This boundary curve is “U”-shaped which means QQ and QSI have a synergistic effect in inhibiting QS^37^. η(QQ) is assumed to be proportional to QQ in this simulation. However, if another enzymatic dynamics such as a Michaelis–Menten equation^38^ were used, η’ (QQ) > 0 and η″(QQ) ≤ 0 are satisfied. If we change the η(QQ) axis to QQ in Fig. 2B, the curve will still be “U”-shaped and the conclusions of the simulation will remain the same.

**Figure 2.**
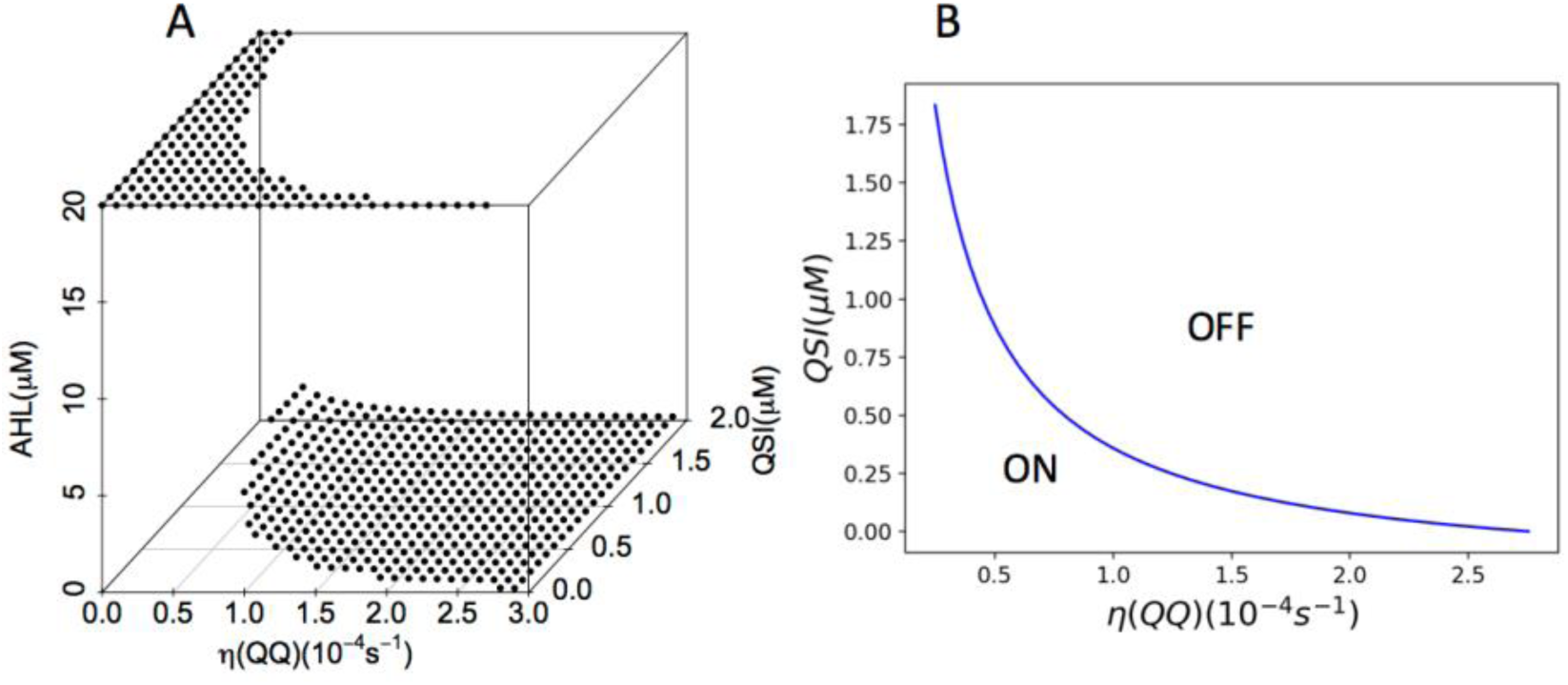
Simulation QS states to QQ and QSI. (A) 3D stationary AHL concentration to η(QQ) and QSI, (B) 2D map of QS on and off states to η(QQ) and QSI.

### Synergistic effects on QS bioreporter strains

To validate the mathematical modeling results, the synergistic effects of AiiA enzyme and G1 were tested using the *P. aeruginosa* QS bioreporter strain PAO1-*lasB-gfp*^39^. The elastase encoding *lasB* gene is controlled by LasR and any induction in the fluorescence signals would indicate the presence of 3-oxo-C12-HSL^40^. The compounds were tested at different concentration gradient to generate dose-dependent curves and calculate the IC_50_ values, which represent half of the concentration required to inhibit the gene expression. Most importantly, in support of our non-growth inhibitory antimicrobial principle^1^, neither compounds nor enzymes affect the growth rate of the bacteria (Supplementary Figure S1). The reduction of the GFP output was indeed due to the effect of compounds in reducing expression of the QS controlled *lasB*-*gfp* gene. The growth measured as OD_600_ was used as control of our non-growth inhibitory concept.

Both G1 and AiiA inhibited *lasB-gfp* expression in dose-dependent manner with IC_50_ values of 13.33±2.37 µM and 4.58±1.05 µM respectively. Promising results were obtained in combinational therapy of AiiA and G1, where the IC_50_ values were significantly reduced to low micromolar range. IC_50_ values calculated for G1 when combined with 32 µg/mL of AiiA was 6.98±1.98 nM and 58.65±19 nM with 16 µg/mL of AiiA. The *lasB-gfp* readings were much lower in the combination treatment as compared to single treatments (Fig. 3).

**Figure 3.**
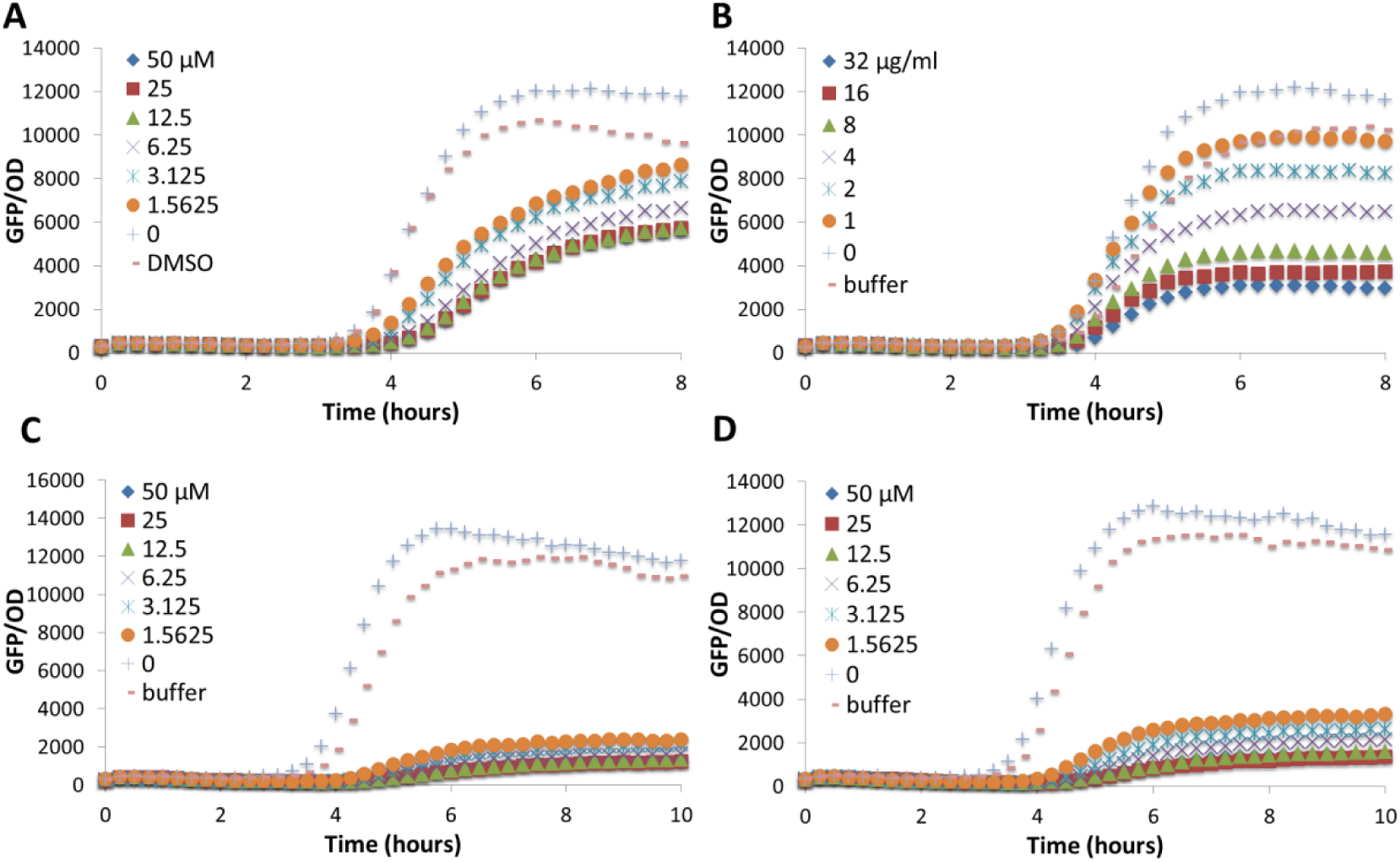
Dose-dependent curves of compounds with QS reporter strain PAO1-*lasB-gfp.* 587 (A) G1, (B) AiiA, (C) G1 and AiiA at 32 µg/mL, and (D) G1 and AiiA at 16 µg/mL. 588 Experiments were done in triplicate manner, only representative data are shown.

Next, we investigated if the synergistic effect would also affect the PQS system, the third intercellular signaling mechanism of *P. aeruginosa* that regulates numerous virulence factors, including those involved in iron scavenging and apoptosis of host cells^41,42^. PQS is under positive regulation of LasR and negative regulation of RhlR^41,43^. PQS has also been detected in the lung of cystic fibrosis patients^44^ and reported to suppress host innate immune responses through nuclear factor-κB pathway^45^.

For this experiment, we tested the compounds against *pqsA-gfp* reporter fusion. The biosynthesis of PQS and other classes of alkyl quinolones requires genes encoded by the *pqsABCDE* and *phnAB* operons^46^. Interestingly, the AiiA didn’t show much inhibition effects on *pqsA-gfp* with IC_50_ values calculated to be 15.58±0.17 µM. Combination treatment between G1 and AiiA significantly reduced the *pqsA-gfp* expression to IC_50_ values of 0.63±0.06 µM (Fig. 4).

**Figure 4.**
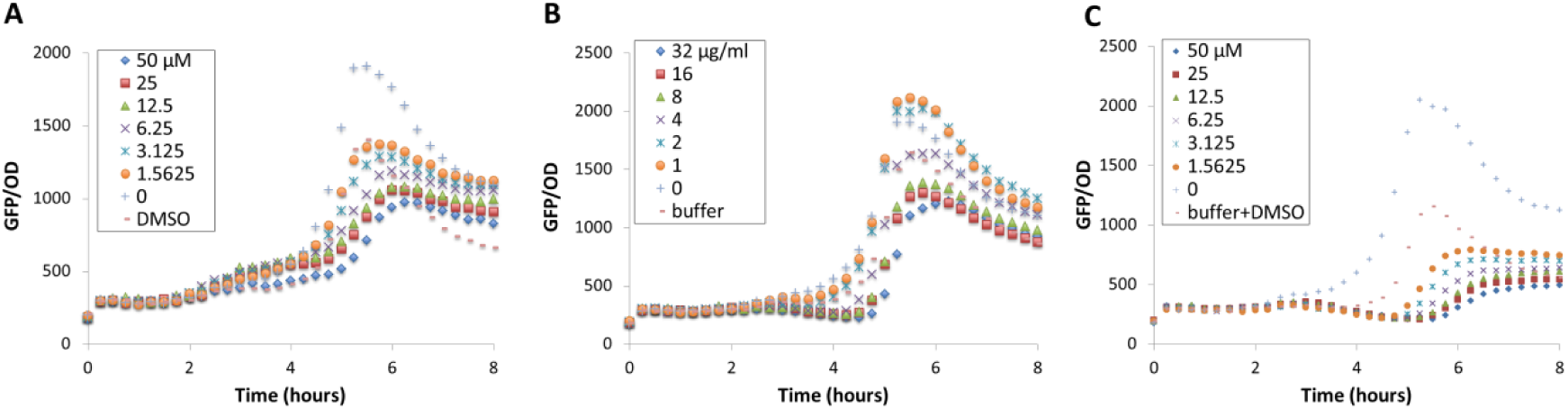
Dose-dependent curves of compounds with QS reporter strain PAO1-*pqsA-gfp.* (A) G1, (B) AiiA, (C) G1 and AiiA at 32 µg/mL. Experiments were done in triplicate manner, only representative data are shown.

### G1 has different affinity to the LasR and RhlR proteins

We next examined the synergistic effects on the *rhl* system, which regulates many QS-dependent virulence factors once activated upon formation of RhlR-C4-HSL^10,47^. The AiiA has been experimentally shown to degrade C4-HSL^48^. Our previous experiments also showed that G1 was able to inhibit *rhl* system more effectively in *P. aeruginosa lasR* mutant but not the *rhl* system in the PAO1 wildtype^31^. We thus hypothesized that G1 has different binding affinity to LasR than RhlR in the PAO1 wildtype and its intracellular concentration is not high enough to repress both LasR and RhlR simultaneously.

To test this hypothesis, we examined the competitive binding efficacy of G1 with 3-oxo-C12-HSL and C4-HSL using a QS deficient *P. aeruginosa* Δ*lasI*Δ*rhlI* double mutant which can respond to the addition of exogenous AHLs (3-oxo-C12-HSL and C4-HSL respectively). In this setting, only one QS system is activated at one time. The reporter strains showed dose-dependent curves when supplemented with different concentration of 3-oxo-C12-HSL and C4-HSL (Fig. 5A and 5C). When 50 µM of G1 was added together with 3-oxo-C12-HSL, we only observed reduction in *lasB-gfp* as compared to the control when concentration of 3-oxo-C12-HSL is below 1.25 µM. However, G1 was able to reduce *rhlA-gfp* expression with all the tested C4-HSL concentrations (up to 10 µM) (Fig. 5B and 5D). Thus we suggest that G1 has a higher affinity to the RhlR than the LasR. However because most of the intracellular G1 was consumed due to LasR abundance, hence its effect to inhibit *rhl* QS in the PAO1 wildtype was abolished due to the earlier induction of *las* QS than the *rhl* QS during growth.

**Figure 5.**
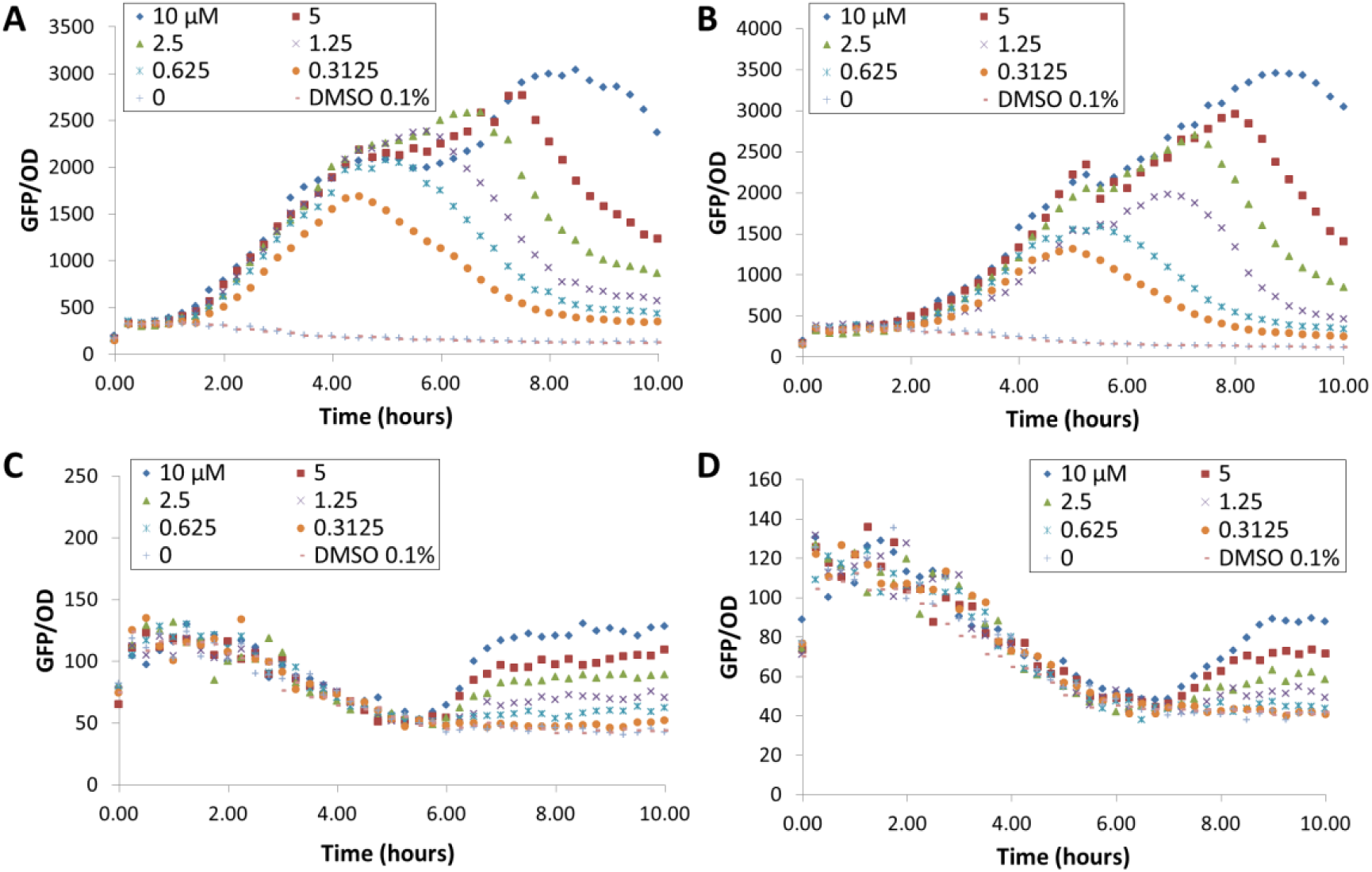
Dose-dependent curves of QS deficient Δ*lasI*Δ*rhlI* double mutant harboring *lasB-gfp* (top) and *rhlA-gfp* (bottom) supplemented with (A) 3-oxo-C12-HSL, (B) 3-oxo-C12-HSL with G1 50 µM, (C) C4-HSL, and (D) C4-HSL with G1 50 µM. Experiments were done in triplicate manner, only representative data are shown.

### AiiA enhances inhibition of G1 on *rhl* QS system in *P. aeruginosa*

Since AiiA has a strong synergy with G1 in inhibiting *las* system, we hypothesized that AiiA is able to have a synergistic effect with G1 in inhibiting the *rhl* system due to the fact that low abundance of LasR protein will ‘consume’ less amounts of G1 in the presence of AiiA, hence more G1 molecules could bind with RhlR. For this experiment, we used PAO1-*rhlA-gfp* bioreporter strain to study the synergistic effects of both compounds. The *rhlA* is the first gene of the *rhlAB* operon that codes for the rhamnolipid biosynthesis^47^. We observed similar findings where the *rhlA-gfp* activity was highly suppressed in the combination treatment (Fig. 6A-C). IC_50_ values calculated for the combination treatment between G1 and AiiA for the *rhlA-gfp* expression is 17.7±1.4 nM, much lower than the single treatment of QSI and QQ (IC_50_ for G1 = 3.65 ±0.95 µM and AiiA = 17.79 ±1.77 µM).

**Figure 6.**
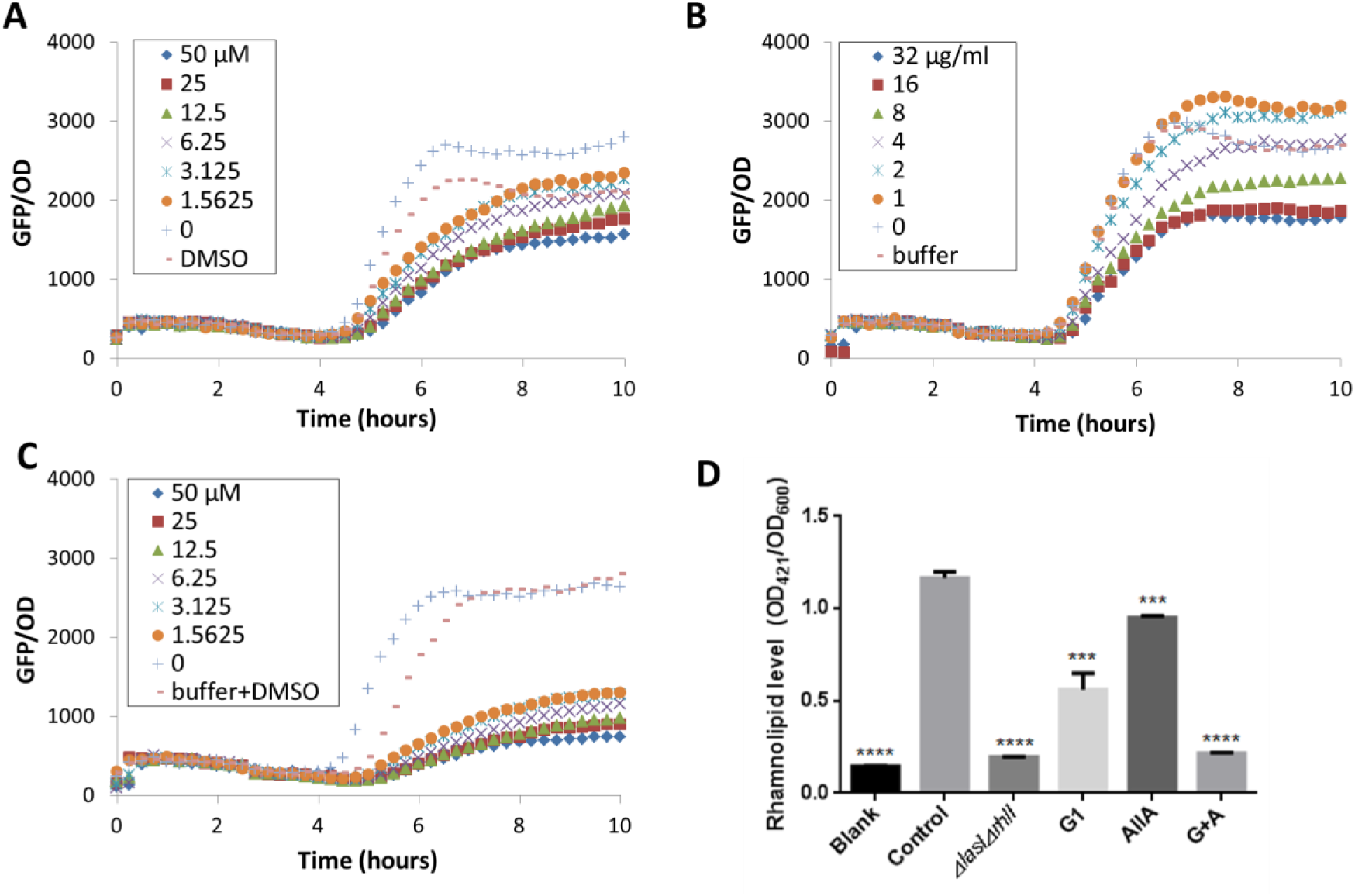
Effects of QQ and QSI on the *rhl* system. Dose-dependent curves of compounds with QS reporter strain PAO1-*rhlA-gfp* (A) G1, (B) AiiA, (C) G1 and AiiA at 32 µg/mL. (D) Effects on rhamnolipid production when tested at final concentration of 50 µM (for G1), and 32 µg/mL (for AiiA). Same amount of DMSO and buffer were used as positive control. PAO1 Δ*lasI*Δ*rhlI* was used as negative control. Experiments were done in triplicate manner. Error bars are means ± SDs. *** = p < 0.001. **** = p < 0.0001, Student’s t test.

A strong synergistic effect was observed from combination of AiiA and G1, thus we were interested to investigate if the synergistic effects could reduce the virulence of *P. aeruginosa*. We decided to test the rhamnolipid production, as it is one of the key QS regulated virulent factors in the early stages of infection. Rhamnolipid promotes infiltration of respiratory epithelia cells^49^ and promote rapid necrotic killing of polymorphonuclear (PMNs) leukocytes^50^. Rhamnolipid is also critical in each stage of biofilm formation and contribute to the structure of biofilms^51,52^.

In the rhamnolipid assay, overnight culture of PAO1 was adjusted to OD_600_ 0.01 and grown in the presence of AiiA, G1 and combination of AiiA and G1 for 18 hours. The rhamnolipid was then extracted and quantified using the orcinol assay^53^. Treatment with AiiA alone didn’t fully decrease rhamnolipid production. However, when combined with G1, the rhamnolipid production was almost diminished to similar level of QS defective Δ*lasI*Δ*rhlI* mutant (Fig. 6D). The findings correlate well with the results obtained from inhibition of *rhlA-gfp* bioreporter strain. The experimental results showed promising application of QQ and QSI in reducing virulence factors associated with host infection.

## Discussion

Over the years, the emergence of multidrug-resistant bacteria and shortage of new antibiotics have been seen as critical issues and greatest threat to human health. Antivirulence approach has been long considered as alternatives in controlling the pathogenicity and reducing the resistance development^1^. In this study, we first reported the synergistic activities of quorum quenching (QQ) enzyme and quorum sensing (QS) inhibitors in inhibiting *P. aeruginosa* LasR/I circuit, one of the important QS regulators. The two classes of compounds intercept QS in different mechanisms, and thus it is interesting to study the synergistic effects. Here, we use mathematical modeling of *P. aeruginosa* LasR/I QS network in batch culture to study whether QQ and QSI have synergistic, antagonist, or additive effects in quenching QS.

Simulation results show that very large η(QQ) or QSI concentration was needed to inhibit QS. When combined, we observed strong synergistic effects between QQ and QSI. Interestingly, switching of QS circuit in the simulations was not observed in experiments. This might be due to simplification of the mathematical model, which assumes every cell is homogeneous and synchronized. In the single-cell study of QS signaling in *V. fischeri,* switching behaviour was observed while tracking single cells but the population level fluorescence was a graded response^34^. This could also be the case of lactose utilization network, where switching was observed in single cell but not at population level^54^.

The experimental results showed promising application of QQ and QSI based on different bioreporter assays. The AHL-dependent QS system has been an attractive target to control bacterial pathogenicity as it controls wide range of virulence gene expression. The AiiA enzyme has been reported to show high specificity and preference towards different signal molecules (acyl chain length and substitution)^48^ and demonstrated to reduce the concentration of 3-oxo-C12-HSL based on our HPLC analysis (Supplementary Figure S2). In some cases, the degradation of QS signal alone is not sufficient to completely diminish and block the QS activities^30^. AiiA could abolish and effectively quench the AHL signal molecules, however it was surprising to see its much lesser effect on the PQS system, which was also under *las* regulation. Combination treatment with G1 resulted in significant reduction of *lasB, pqsA,* and *rhlA-gfp* expression as compared to single treatment of both AiiA and G1. Our work demonstrated that combining two classes of QSI and QQ could provide multiple points of attacks and efficient blockade of QS-mediated signaling pathways.

Although LasR regulator has long been considered essential for full virulence of *P. aeruginosa*^*55*^, loss of function *lasR* mutants occur frequently in the natural environment ^56^ and also cystis fibrosis patients^57^ and individuals suffering from pneumonia and wound infections^58^. In the *lasR* mutants, the QS-regulated virulence factors continue to be expressed. There has also been reports that the *rhl* system could override the hierarchy of QS network in a non-functional *las* system^59^. Recent studies also showed that RhlR plays critical roles as QS regulator using *Drosophila melanogaster* oral infection model^60^ and controls pathogenesis and biofilm development^61^.

In our previous study, G1 has been shown to interact and compete with AHL to inhibit LasR in *P. aeruginosa*^31^. In *P. aeruginosa* PAO1 strain, where both *las* and *rhl* circuits exist, G1 could only inhibit *las* system but cannot inhibit *rhl* system. However, when *lasR* was mutated, G1 can effectively inhibit *rhl* system. In our competitive binding assay using bioreporter strains, we demonstrated that QSIs might have different affinity to the QS receptor proteins. G1 can inhibit *lasB-gfp* expression only when concentration of 3-oxo-C12-HSL is smaller than 1.25 µM. However, the inhibition effect of G1 to *rhlA-gfp* was still significant even when the concentration of C4-HSL is 10 µM. We also observed the transcription rates of *las* system is activated first than that of *rhl.* The results explain that despite of its higher binding affinity to RhlR, G1 would still bind to LasR because the *las* system is activated first in PAO1. But in the case of *lasR* mutant, G1 has higher competitiveness to C4-BHL and could inhibit *rhl* system effectively.

The results also suggested why AiiA could enhance the inhibition effects of G1 on *rhl* system, as shown by our experimental data. Assuming intracellular concentration of G1 is constant, G1 would have to competitively bind to both LasR and RhlR. In this case, addition of AiiA to quench both 3-oxo-C12-HSL and C4-HSL resulted in lower abundance of LasR and therefore G1 can specifically bind to inhibit RhlR. Thus combination of QQ and QSIs might not only enhanced the efficacy of QSIs, but also expand the targeting systems of QSIs as many bacteria have more than one *lux* QS systems.

In conclusion, the synergistic effects of QQ enzyme and QSI compound have been demonstrated *in vitro* in this study. Mathematical modeling showed enhanced QS inhibiting effects on AHL concentration when QQ enzyme (AiiA) and QSI (G1) were applied together. We have also provided better understanding and elucidated QS network interaction with G1 in this work. The implication of our study represents a novel approach of utilizing QS-interfering compounds to impede virulence and block pathogenesis. Future work is aiming to evaluate the effectiveness of the combined treatments *in vivo.*

## Materials and Methods

### General information

All chemicals were purchased from Sigma Aldrich and used without further purification. G1 was purchased from TimTec LLC (Newark, DE). Bacteria were grown in Luria-Bertani (LB) broth (1% tryptone, 0.5% yeast extract, and 0.5% NaCl). Media used for biological assay was ABTGC (AB minimal medium supplemented with 0.2% glucose and 0.2% casamino acids)^62^. Bacterial strains used in this study are shown in Table 1.

**Table 1.**
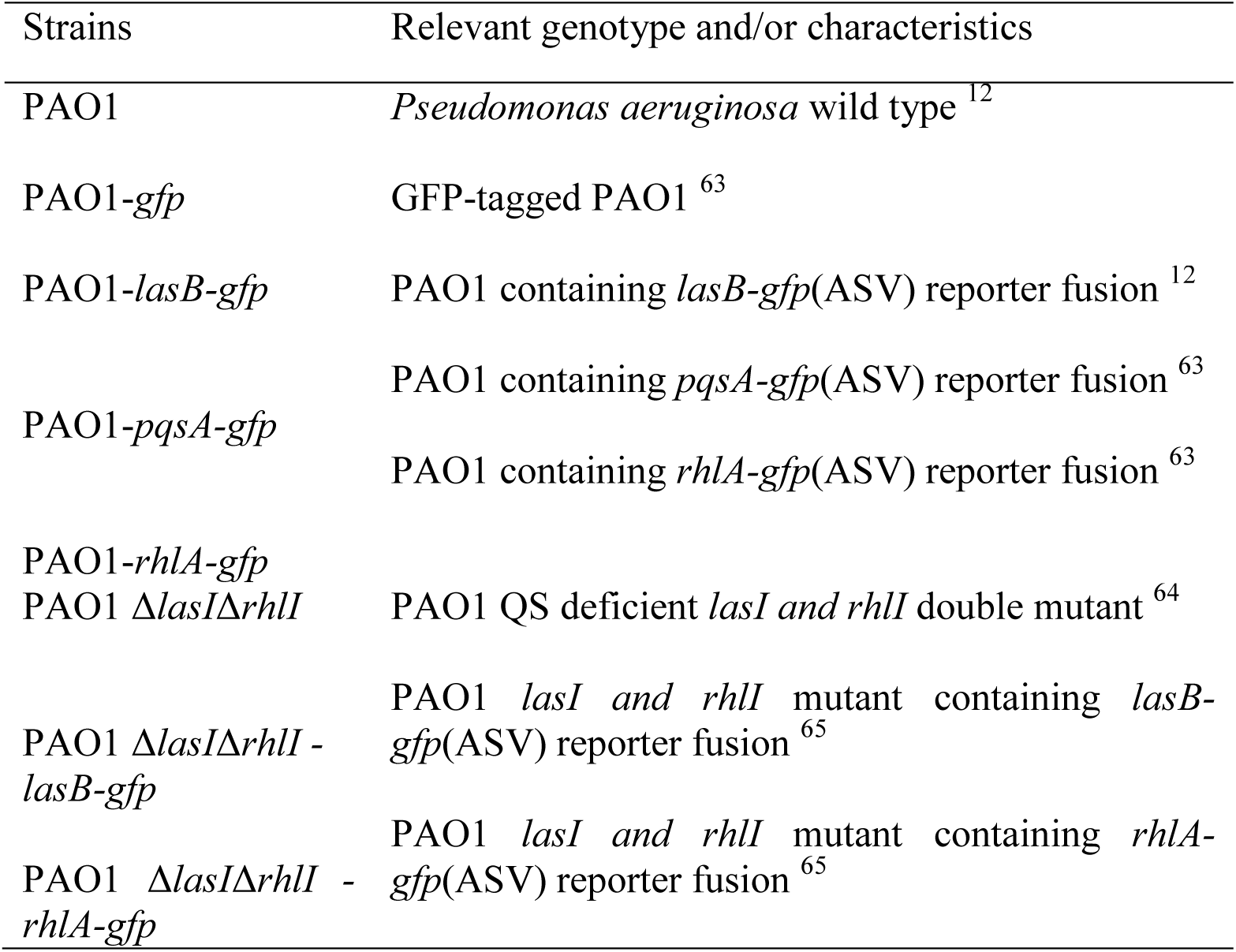
Bacterial strains used in this study.

| Strains | Relevant genotype and/or characteristics |
| --- | --- |
| PAO1 | *Pseudomonas aeruginosa* wild type<sup>12</sup> |
| PAO1-*gfp* | GFP-tagged PAO1<sup>63</sup> |
| PAO1-*lasB-gfp* | PAO1 containing *lasB-gfp*(ASV) reporter fusion<sup>12</sup> |
| PAO1-*pqsA-gfp* | PAO1 containing *pqsA-gfp*(ASV) reporter fusion<sup>63</sup> |
| PAO1-*rhlA-gfp* | PAO1 containing *rhlA-gfp*(ASV) reporter fusion<sup>63</sup> |
| PAO1 Δ*lasI*Δ*rhlI* | PAO1 QS deficient *lasI* and *rhlI* double mutant<sup>64</sup> |
| PAO1 Δ*lasI*Δ*rhlI* - *lasB-gfp* | PAO1 *lasI* and *rhlI* mutant containing *lasB-gfp*(ASV) reporter fusion<sup>65</sup> |
| PAO1 Δ*lasI*Δ*rhlI* - *rhlA-gfp* | PAO1 *lasI* and *rhlI* mutant containing *rhlA-gfp*(ASV) reporter fusion<sup>65</sup> |

### Expression and purification of QQ enzyme AiiA

The gene coding for AiiA was cloned into pET-47b(+) vector (BamHI-HindIII sites). The expression vector was transformed into *E. coli* BL21(DE3) competent cells (New England Biolabs, USA). The cells were grown in 2L of LB media supplied with 35 mg/L Kanamycin at 37°C. The expression of the protein was induced with 0.5 mM of Isopropyl β-D-1-294 thiogalactopyranoside (IPTG) when OD_600_ reached 0.6. The cells were grown for overnight at 18°C. The harvested cells were resuspended in 50 mL of lysis buffer (50 mM Tris-HCl, pH 8.0, 150 mM NaCl, 0.05% (v/v) CHAPS, 10 % (v/v) glycerol) and lysed by passing the homogenized cells through an Emulsiflex-C3 (Avestin, USA) high-pressure apparatus at 15,000 psi three times. The cell lysate was centrifuged at 25,000 g for 25 min. The supernatant was then applied to Ni-NTA gravity column (Bio-rad) equilibrated with lysis buffer. After extensive washing with lysis buffer, the bound proteins were eluted with lysis buffer containing increasing concentration of imidazole (0 - 300 mM). The eluted fractions were analyzed with 15% of SDS-PAGE and fractions containing the desired protein were pooled and dialyzed against lysis buffer. The final concentration of the protein was measured using Bradford Assay.

### Enzymatic assay of AiiA

The 3-oxo-C12-HSL hydrolysis activity of AiiA was tested with 500 μM 3-oxo-C12-HSL, 30 μM AiiA in reaction buffer (20 mM Tris-HCl, 150 mM NaCl, pH 8.0). After 30 min of reaction at 30°C, the reaction mixture was monitored at 215 nm by analytic C18 reverse phase HPLC column (Jupitar, 5μ, 300Å, 250x4.6 mm) with a flow rate of 0.5 mL/min (Gradient: 0-100% buffer B (90% acetonitrile, 10% H2O, 0.05% TFA) in buffer A (100% H2O, 0.05% TFA) for 50 min).

### Model of LasR/I circuit

The models in this study simulated batch cultures according to the experimental setup. The LasR/I QS circuit of *P. aeruginosa* is shown in Fig. 7. QSI binds to LasR similarly as AHL, but in this case only AHL can stabilize the LasR^66^. Since our focus of the present study was whether QQ enzyme and QSI have synergistic effect in inhibiting QS, some complex features in the QS network were simplified to make the model easier to implement and reduce computational cost. For instance, the interactions of LasR/I QS circuits with other cellular components, such as the binding of 3-oxo-C12-HSL to RhlR^21^ were not included in the network studied in this work. Both heterogeneity and asynchronization of cells were beyond the scope of the modeling in this work. The final component concentrations of cells in batch culture were adapted in the computation of AHL concentration. As the response time of QS switching is much faster than the time required for culture growth^34^, the final component concentrations can be approximated by the stationary concentrations with the final cell volume fraction **ρ**. The AHL concentration was considered to be homogeneous inside and outside cells due to its large diffusion coefficient^67^. Vfr was assumed to be some large enough constant since it is normally expressed in experimental strains of this study. QSI concentration was taken as a constant considering its relative big value. A maximum concentration of AHL **A**_**max**_ was set in the model to avoid very large concentrations of components caused by the accumulation of stable AHL in batch cultures^68^. The reactions of QS network are shown in Table 2. QQ enzyme and QSI are written as **Q**_**Q**_ and **Q**_**I**_ to avoid confusion when necessary.

**Figure 7.**
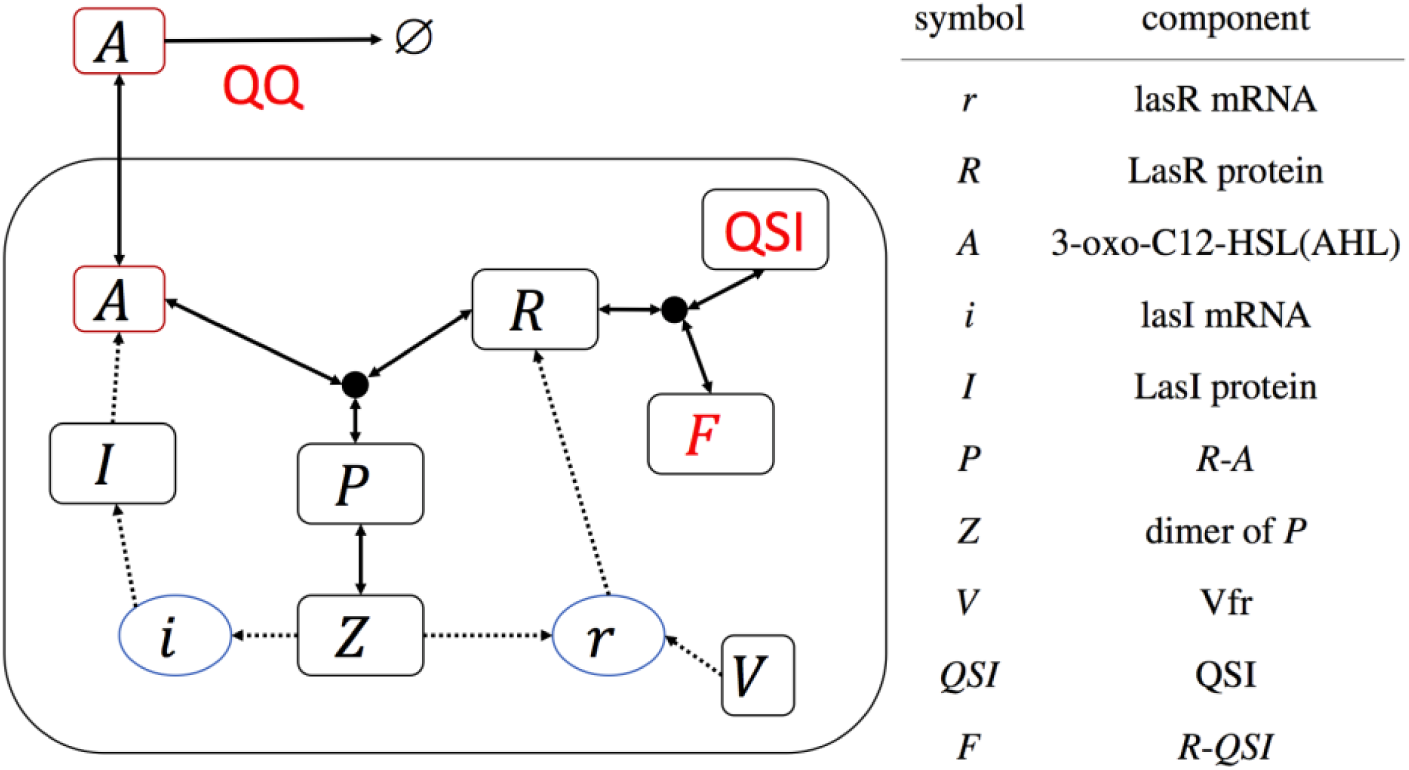
*P.aeruginosa* LasR/I circuit model with QQ and QSI both indicated in red. Dashed lines indicate the reactants still remains after the reactions and solid lines indicate the reactants will disappear after the reactions.

**Table 2.** Biochemical reactions in *P.aeruginosa* LasR/I circuit.

| Reaction | Description | Rate |
| --- | --- | --- |
| $I \rightarrow I + A$ | Production of A | $\frac{V_A I}{K_A + I}$ |
| $A \rightarrow \text{null}$ | Natural decay of A | $d_A A$ |
| $R + A \rightarrow P$ | Combination of R and A | $k_{RA} R A$ |
| $P \rightarrow R + A$ | Dissociation of P | $d_P P$ |
| $R \rightarrow \text{null}$ | Decay of LasR protein | $d_R R$ |
| $r \rightarrow r + R$ | Translation of lasR mRNA | $k_r r$ |
| $\text{null} \rightarrow r$ | Transcription of lasR | $r_0 + V_r Z \left( \frac{1 - e^{-\beta V}}{K_{r1} + Z} + \frac{e^{-\beta V}}{K_{r2} + Z} \right)$ |
| $r \rightarrow \text{null}$ | Decay of lasR mRNA | $d_r r$ |
| $i \rightarrow i + I$ | Translation of lasI mRNA | $k_i i$ |
| $I \rightarrow \text{null}$ | Decay of LasI protein | $d_I I$ |
| $\text{null} \rightarrow i$ | Transcription of lasI | $i_0 + \frac{V_i Z}{K_i + Z}$ |
| $i \rightarrow \text{null}$ | Decay of lasI mRNA | $d_i i$ |
| $2P \rightarrow Z$ | Dimerization of P | $k_Z P^2$ |
| $Z \rightarrow 2P$ | Dissociation of Z | $d_Z Z$ |
| $R + Q_I \rightarrow F$ | Combination of R and QSI | $k_{RQ} R Q_I$ |
| $F \rightarrow R + Q_I$ | Dissociation of F | $d_F F$ |
| $F \rightarrow Q$ | Degradation of R in F | $d_R F$ |
| $A \rightarrow \text{null}$ | Degradation of A by QQ | $\eta(Q_Q) A$ |

The ordinary differential equations of the QS network are listed in equations 1 to 8. Stationary QS components were solved using the steady state condition. When there are multiple stable stationary solutions, the state with smallest concentrations was chosen as the outcome presented in this work. The parameters of *P*. *aeruginosa* LasR/I QS circuit were firstly estimated from reported values in the literature, then optimized to enable the switching behaviour of the QS network observed in experiments (Supplementary Table S1)^34,35^.

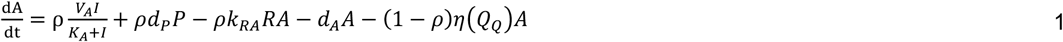

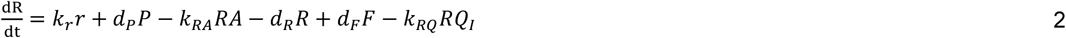

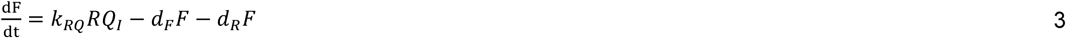

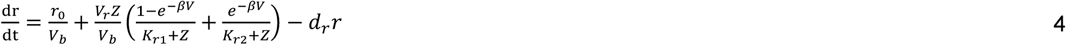

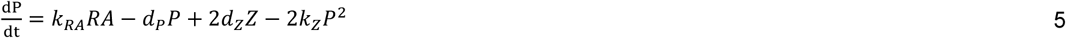

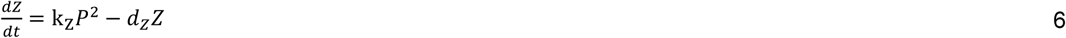

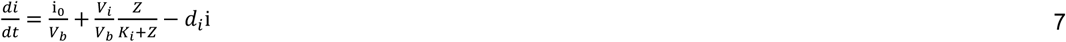

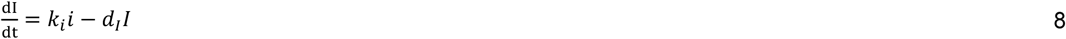

### Reporter gene assay

Stock solution of G1 was prepared by dissolving appropriate amount of chemicals in DMSO to make final concentration of 10 mM, aliquoted into small Appendorf tube and stored at −20°C until further usage. The compound was then dissolved in ABTGC medium to the working concentration and 100 µL of this solution was pipetted into first rows of 96-well microtiter dish (Nunc, Denmark). 2-fold serial dilution was made to the rest of the rows, leaving last two rows empty for blank and solvent control. Next, 50 µL of AiiA diluted in ABTGC media was added into each well. Overnight culture of *P. aeruginosa* reporter strain PAO1-*lasB-gfp* was diluted to optical density at 600 nm (OD_600_) of 0.02 (approximately 2.5 x 10^8^ CFU/mL). 100 µL of the bacterial suspension was added to each wells and the plate was incubated for 18 hours at 37°C. GFP fluorescence (excitation at 485 nm, emission at 535 nm) and OD_600_ readings were recorded every 15 mins using Tecan Infinite 200 Pro plate reader (Tecan Group Ltd, Männedorf, Switzerland). IC^50^ values were calculated using Graphpad Prism 6 software. All assays were done in triplicate manner.

### Rhamnolipid quantification

Rhamnolipid was extracted and quantified using method reported by Koch et al. with modifications^53^. Briefly, overnight culture of *P. aeruginosa* was diluted to OD_600_ 0.01 in ABTGC medium. Into the cultures, compounds were added to appropriate concentration and the cultures were grown for 18 h at 37°C, shaking condition (200 rpm). Supernatants were collected and extracted with diethyl ether twice. The organic fractions were collected and concentrated to give white solids, which were further dissolved in water. 0.19% (w/v) orcinol in 50% H^2^SO^4^ was freshly prepared and 363 added into the water solution. It was then heated at 80°C for 20-30 min to give yellow-orange solution. The solution was allowed to cool at room temperature before measuring the absorbance at 421 nm. The results were normalized with cell density at OD_600_. Experiments were done in triplicate manner.

## Acknowledgements

This research was supported by the National Research Foundation and Ministry of Education Singapore under its Research Centre of Excellence Program (SCELSE) and AcRF Tier 2 (MOE2016-T2-1-010) from Ministry of Education, Singapore.

## Author Contributions

H.B.S., B.W. and L.Y. designed methods and experiments. C.Z., R.Y. and J.F. carried out the laboratory experiments, analyzed the data, and interpreted the results. L.Y. and M.G. discussed analyses, interpretation, and presentation. J.F., C.Z., and L.Y. wrote the paper. All authors have contributed to, seen, and approved the manuscript.

## Additional Information

All data generated or analysed during this study are included in this published article and its Supplementary Information files.

The authors declare no competing financial interests.

